# C1q in non-immune human serum has a non-redundant complement function towards clinical Mycobacterium tuberculosis complex strains

**DOI:** 10.1101/2024.11.01.621467

**Authors:** Mario Alejandro Duque, Maximilian Peter Götz, Emilie Rousseau, Susanne Homolka, Stefan Niemann, Peter Garred, Christoph Hölscher, Kerstin Walter, Anne Rosbjerg

**Author notes:** KW and AR share last authorship.

## Abstract

The *Mycobacterium tuberculosis* complex (MTBC) includes three human-adapted species: *Mycobacterium tuberculosis*, *Mycobacterium africanum*, and *Mycobacterium canettii*. With their genetic diversity and inherent virulence, these bacteria interact with host factors to influence tuberculosis (TB) transmission and pathogenesis. A critical yet poorly understood aspect of the immune response in TB involves the role of the complement system. Our study aimed to assess the general interaction between the complement system and key clinical MTBC strains representing the principal lineages known to vary in virulence. We found that not only mannose binding lectin (MBL) from the lectin pathway, but also C1q from the classical pathway directly recognize mycobacteria. While MBL generally showed a higher level of binding than C1q, the binding interactions varied across different MTBC strains. We further observed that complement activation products C4b, C3b and the terminal complement complex (TCC) were deposited on the surface of MTBC strains. Inhibition of MBL alone did not substantially alter complement activation, whereas C1q inhibition ceased activation for most strains investigated. Exposure to human serum did not impact the viability or growth of the MTBC strains. In conclusion, both the classical and lectin pathways of complement via C1q and MBL are activated by a broad range of lineages of the MTBC. The classical pathway is the main complement activator in non-immune serum across diverse MTBC lineages, with C1q playing a potentially novel role in the immune response in TB.

## Introduction

Tuberculosis (TB) remains a major global health threat. In 2022 a total of 7.5 million people were newly diagnosed with TB, which is the highest recorded number since the WHO began global TB monitoring. It is estimated that 1.3 million patients lost their lives in 2022, thus TB belongs to the most deadly infectious diseases worldwide [1].

The causative microorganisms of TB are bacteria of the *Mycobacterium tuberculosis* complex (MTBC), which have a considerable genomic diversity [2, 3]. Within the human-adapted lineages (L) of the MTBC, different main lineages of *Mycobacterium* (*M.*) *tuberculosis* such as L1, L2, L3, L4 and of *M. africanum* (L5, L6) were described to cause TB in different parts of the world [4]. The phylogenetically “modern” lineages L2, L3 and L4 [5] are globally spread; L2 (East Asian) is distributed in East Asian countries, L3 (Dehli/CAS) appears in East Africa and Central and South Asia and L4 (Euro-American) is widely spread in America, Europe, the Middle East and Africa. Phylogenetically “ancient” strains are geographically more restricted and include L1 (East African Indian) which occurs around the Indian Ocean and the Philippines [2] and the *M. africanum* lineages L5 (West African 1) and L6 (West African 2) that are restricted to Western African countries where they show a prevalence of up to 50% among TB cases [6, 7].

The pathogenomic diversity of MTBC strains can influence the virulence of strains, host-pathogen interactions, the immune response and the disease outcome [8]. Accordingly, we have previously demonstrated that strains from L2 and L4 exhibit enhanced growth in human macrophages and in the mouse model of TB compared to strains of lineages L1, L5 and L6 [9]. Furthermore, we could delineate a conserved core and lineage-specific transcriptome profiles during intracellular survival [10], and describe major differences in the evolution of drug resistance between strains of different MTBC lineages [11, 12]. These findings indicate that genetic variations between MTBC lineages directly affect the interaction of mycobacteria and host sensors and are thus causative for different virulence characteristics of these strains.

The complement system constitutes a rapid and efficient sensing system of the host and acts as a first line of defense in detecting and removing pathogens. The three following pathways activate the complement cascade: The classical pathway (CP), which is typically induced by C1q binding to antigen-attached antibodies. The lectin pathway (LP), which is induced by collectins including mannose-binding lectin (MBL), CL-10 and CL-11 and ficolins 1, 2 and 3. The alternative pathway (AP), which is spontaneously activated at a low rate and, moreover, acts as an amplifier of the classical and lectin pathways. The main effector functions of activated complement are the opsonization of target cells by proteins (e.g. C3b), lysis via formation of the terminal complement complex (TCC), and inflammation driven by anaphylatoxins, in particular C3a and C5a [13].

Even though MTBC strains and humans share a long co-evolutionary history [3, 4], the relevance of complement in TB has been inadequately addressed so far. A theory, based on clinical findings, has been put forward that the genetic landscape of the complement system and in particular of the lectin pathway has been shaped by the challenge of mycobacterial infections [14, 15]. Nevertheless, most findings regarding the complement system and mycobacteria are based on studies using a limited number of strains, as well as clinically irrelevant strains such as the lab-adapted strains H37Rv [16–18], Erdman [16, 19, 20], *M. bovis* BCG [17, 18, 21–24], or *M. avium* [25]. Studies using only lab-adapted strains will likely skew the picture and disregard lineage differences e.g. with regard to their virulence. Remarkably, by investigating clinical strains of the MTBC, we demonstrated previously that MBL binds to strains of *M. africanum* L6 to a higher extent than to strains of L4, and that a specific MBL2 genetic variant confers protection against TB caused by *M. africanum* [26].

Overall, the relevance of the complement system for TB, in particular for host-pathogen interaction, is only poorly understood, primarily due to a lack of complement activation analyses using MTBC clinical strains. Therefore, this study aims to investigate the interaction of the complement system with clinical strains belonging to 6 different MTBC lineages. Our study will shed light on important aspects of the immune response to MTBC infection and may spark new impetus for further research on complement in TB where relevant MTBC clinical strains must be given more consideration.

## Results

### Genetic diversity and phylogeny of clinical strains

For a broad assessment of interactions between the complement system and MTBC pathogens, a total of 22 clinical strains covering 6 main phylogenetic lineages of the MTBC that are either globally distributed (L2, L3 and L4) or geographically restricted (L1, L5, and L6) were investigated. Whole-genome sequencing data were used to calculate a phylogenetic maximum likelihood tree based on a total of 7476 informative sites differentiating any of the 22 strains investigated. The phylogenetic tree confirms the distinct grouping of the *M. tuberculosis* strains in four groups: L1 (1797/03, 4858/08, 8316/09), L2 (12594/02, 3936/02, 178/03), L3 (2383/12, 38/12, 7660/06) and L4. For L4, the laboratory H37Rv reference strain TMC 102 (H37Rv; ATCC, 27294) and strains of two sub-clusters were investigated that are either globally distributed L4 generalists (L4G) (2336/02, 4130/02, 9532/03) or geographically restricted L4 specialists (L4S) (5390/02, 5400/02, 1417/02) [26, 27]. The other main groups contain *M. africanum* strains of L5 (10494/01, 1449/02, 10415/02, 5434/02) and L6 (10476/01 and 10514/01) (**Fig 1**).

**Figure 1.**
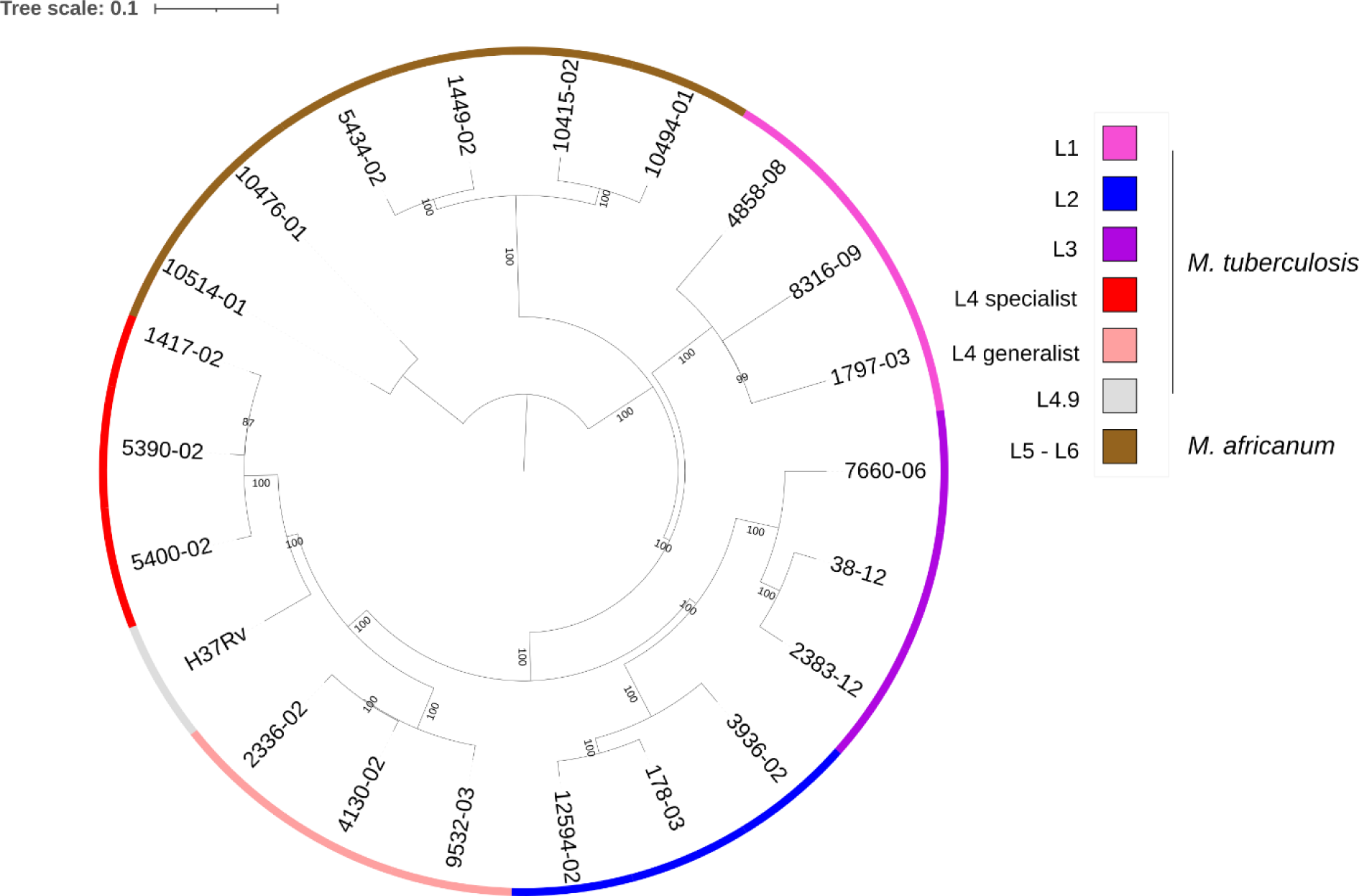
Phylogenetic tree of *Mycobacterium tuberculosis* complex strains used in this study. A midpoint-rooted maximum likelihood tree was calculated based on a total of 7476 concatenated single nucleotide polymorphism (SNPs) derived from whole genome sequencing data of the 22 MTBC strains investigated in the study. In the tree, calculated under the K3Pu+F substitution model, bootstrap values of the main branches are displayed.

### Binding of complement proteins to clinical strains of the MTBC

In an initial experiment, a broad spectrum of complement pattern recognition molecules (PRM), such as ficolin-1, -2, -3, MBL, collectin-11 (CL-11) and C1q, were screened for their ability to bind to clinical strains of the 6 MTBC lineages (L1-L6) and H37Rv. Because MBL-associated serine protease 1 (MASP-1) and -3 were previously shown to also bind pathogens directly [28], these two proteases of the lectin pathway were also included. The binding of purified (p) or recombinant (r) complement proteins was detected by flow cytometry after incubation with specific antibodies against the different components (for gating strategy, refer to **Fig S1**). The comprehensive screening of complement PRMs and MASPs revealed that especially rMBL, rficolin-2 and pC1q bind to the investigated strains of the MTBC (**Fig 2**). MBL recognized all strains, with some variation between the strains of different lineages; the lowest binding was observed for L2 (178/03) and L6 (10476/01) (**Fig 2 A**). Ficolin-2 was mostly detected on strains of lineages L1, L3, L4S and H37Rv (**Fig 2 B**). Interestingly, we also observed a direct binding of C1q on clinical strains of the MTBC. The highest levels of C1q binding were detected on strains of lineages L1, L3, L4S, and H37Rv (**Fig 2 C**). The remaining PRMs and MASPs tested (rficolin-1, rficolin-3, rMASP-1, rMASP-3 and rCL-11) only recognized some lineages of the MTBC to a minor extent and were not further investigated (**Fig 2 D, E, F, G, H**).

**Figure 2.**
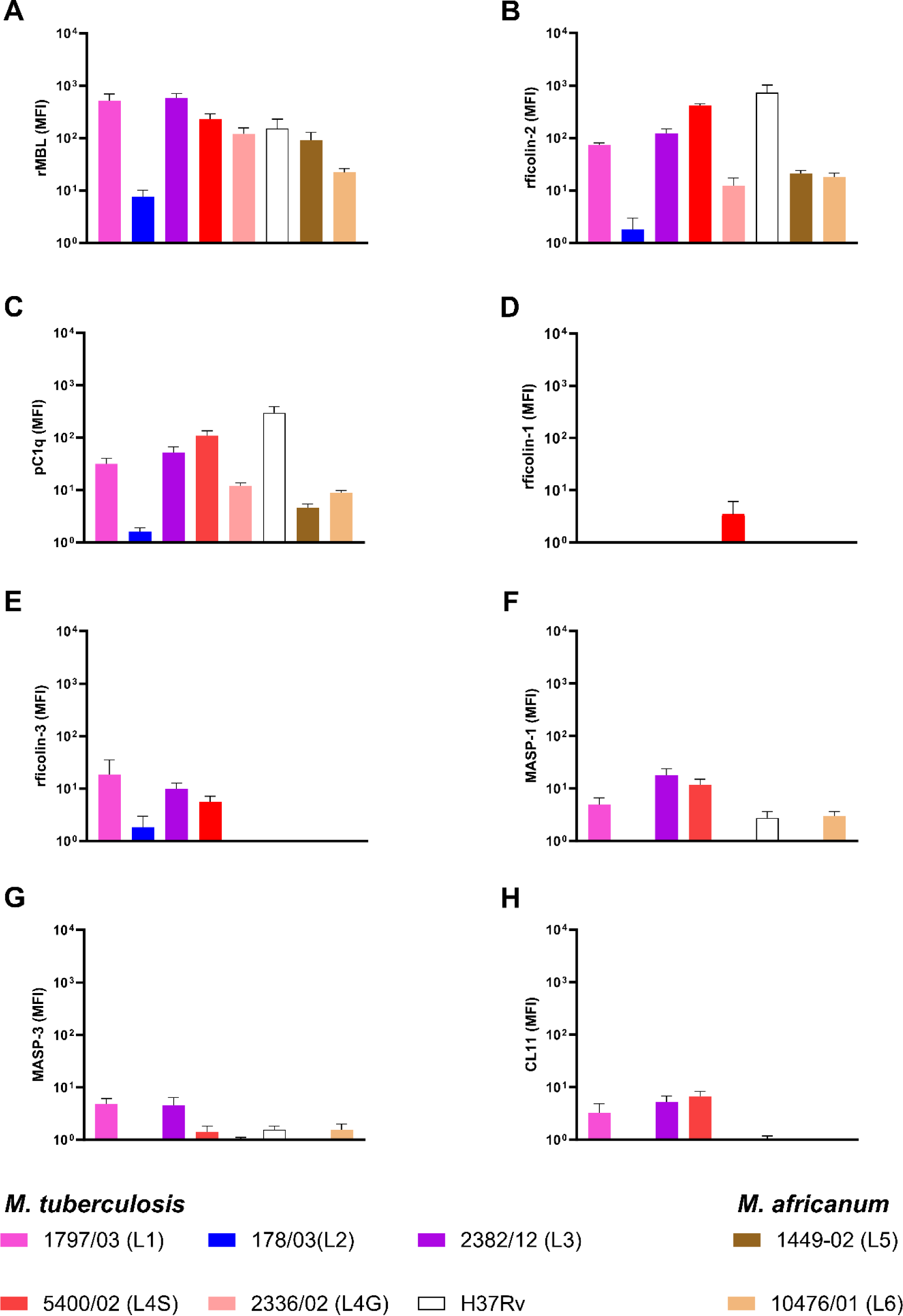
Differential binding of complement components to strains of the *Mycobacterium tuberculosis* complex. Irradiated strains from the MTBC lineages L1, L2, L3, L4S (specialist), L4G (generalist), L5, L6 and H37Rv were incubated with recombinant (r) MBL (**A**), rficolin-2 (**B**), purified (p) C1q (**C**), rficolin-1 (**D**), rficolin-3 (**E**), rMASP-1 (**F**), rMASP-3 (**G**), and rCL-11 (**H**). Complement binding was measured by flow cytometry using specific antibodies and fluorophore-labelled secondary antibodies. Negative controls were processed identically but in the absence of PRM or MASPs. MFI levels indicate the binding after background subtraction and the data represent the means of at least three independent experiments ± SEM.

### Differential binding of MBL and C1q to MTBC strains

In order to confirm and further validate our screening results, we extended the number of clinical strains per lineage and focused on C1q and MBL, as ficolin-2 did not show binding in normal human serum (NHS) (**Fig S2**). The comparison of MBL binding between strains of different MTBC lineages revealed that rMBL bound to all tested strains from lineages L1, L2, L3, L4, L5, L6 and H37Rv, with the lowest level of binding on 178/03 (L2) (**Fig 3A**). By increasing the number of clinical strains, we could observe some variations for pC1q binding to strains of different MTBC lineages. One strain from L2 and one strain from L5 appeared not to bind C1q at all, while both high and low C1q binding was observed within the linages L3 and L4S. Within L1 and L4G, the respective strains bound pC1q to a similar extent. A relatively low level of pC1q binding was detected for all *M. africanum* strains of L5 and L6 (**Fig 3 B**). Hence, by investigating several strains of each lineage we confirmed that MBL binds to strains of all lineages to a higher extent than C1q. Furthermore, we observed a differential binding of these PRMs to strains of MTBC lineages, as *M. africanum* strains (L5 and L6) were recognized by rMBL, whereas pC1q binding was barely detectable.

**Figure 3.**
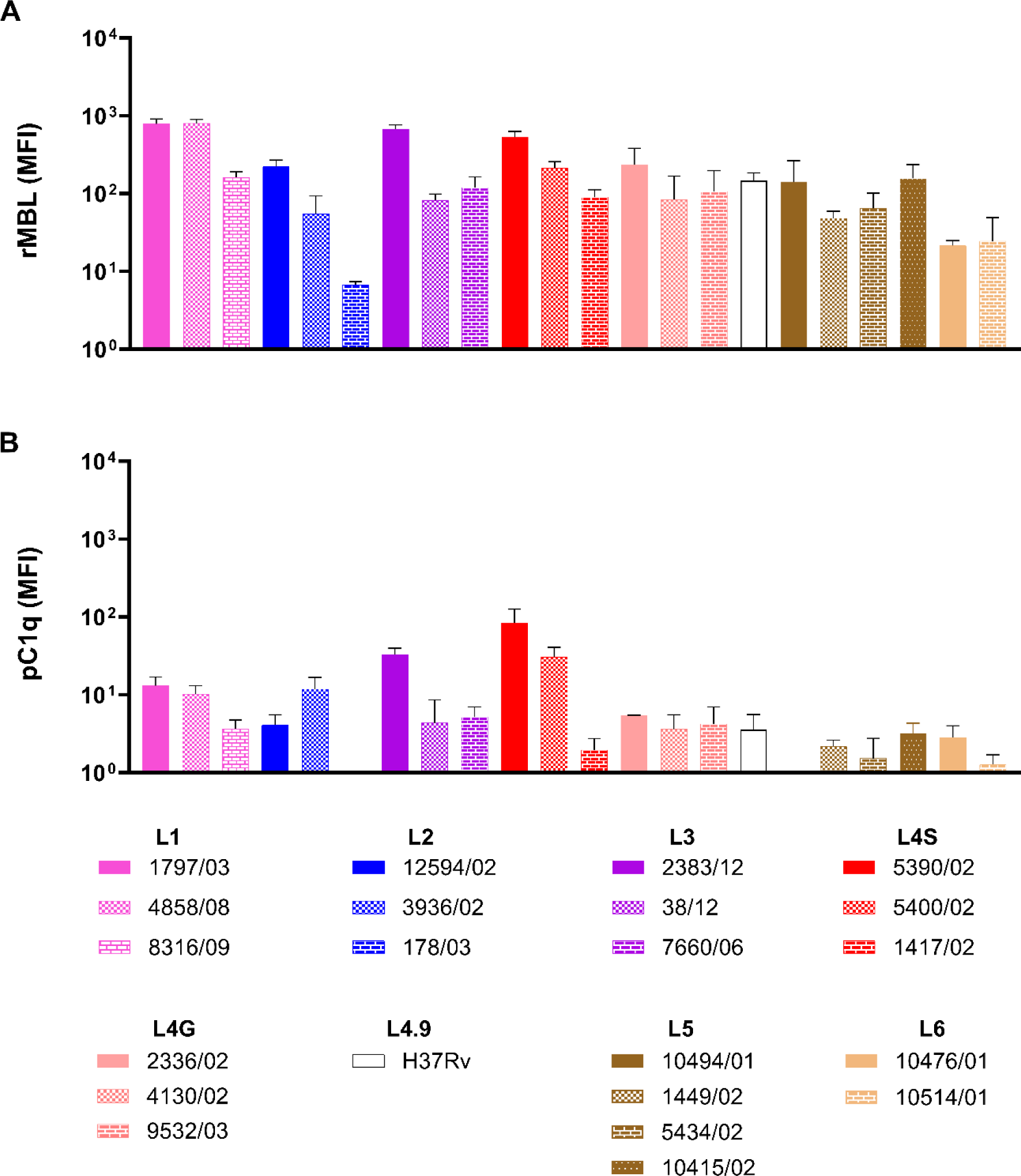
Binding of MBL and C1q to a broad range of clinical strains of different *Mycobacterium tuberculosis* complex strains. Twenty-one irradiated clinical strains covering the *M. tuberculosis* lineages L1, L2, L3, L4S (specialist), L4G (generalist), *M. africanum* lineages L5 and L6 and H37Rv were incubated with recombinant (r)MBL **(A)**, or purified (p)C1q **(B)**. Complement binding was measured by flow cytometry using specific antibodies and fluorophore-labelled secondary antibodies. Negative controls were processed identically but in the absence of rMBL and pC1q. MFI levels indicate the binding after background subtraction and the data represent the means of at least three independent experiments ± SEM.

### Complement activation by main lineages of the MTBC via the CP and the LP

After observing the binding of MBL and C1q to main strains of different MTBC lineages, we next investigated the activation of the complement system and determined the contribution of the different pathways. Since these experiments required a high number of different conditions to be analyzed in parallel for each sample, the number of strains was reduced to one representative clinical isolate per lineage. Complement activation was measured by C3b and TCC deposition on the surface of mycobacteria which was analyzed by flow cytometry. The effect of the CP and the LP was examined by incubating the mycobacteria in 2% NHS (BCG-negative) to reduce the AP activity [29]. The contribution of the respective pathways was investigated by the addition of an anti-MBL (3F8), anti-C1q (CLB/C1q85) or C3 inhibitor (Compstatin, CP40) (**Fig 4**). All *M. tuberculosis* and *M. africanum* strains investigated activated the complement cascade with some variations between the lineages. Interestingly, C3b and TCC deposition were also observed in the presence of the MBL inhibitor indicating that complement is still activated in the absence of the lectin pathway activation for all strains analyzed (**Fig 4**). In contrast, inhibition of C1q strongly reduced C3b and TCC deposition for most strains investigated, suggesting a non-redundant function of this PRM in the activation of the complement cascade. Despite the seemingly insignificant role of MBL regarding complement activation, the simultaneous inhibition of MBL and C1q fully eliminated complement activity. This suggests that MBL is the only LP PRM that activates the complement cascade to a substantial degree even though rficolin-2 was shown to bind strains of the MTBC (**Fig 2B**). As expected, the C3 inhibitor compstatin completely abolished C3b and TCC deposition and incubation with the respective mock inhibitors did not reduce complement activation (**Fig S3**). These results demonstrate that the complement cascade is activated by strains of different MTBC lineages such as *M. tuberculosis* and *M. africanum* strains. Furthermore, complement activation in non-immune serum was mainly driven by the CP, with MBL only playing a minor role.

**Figure 4.**
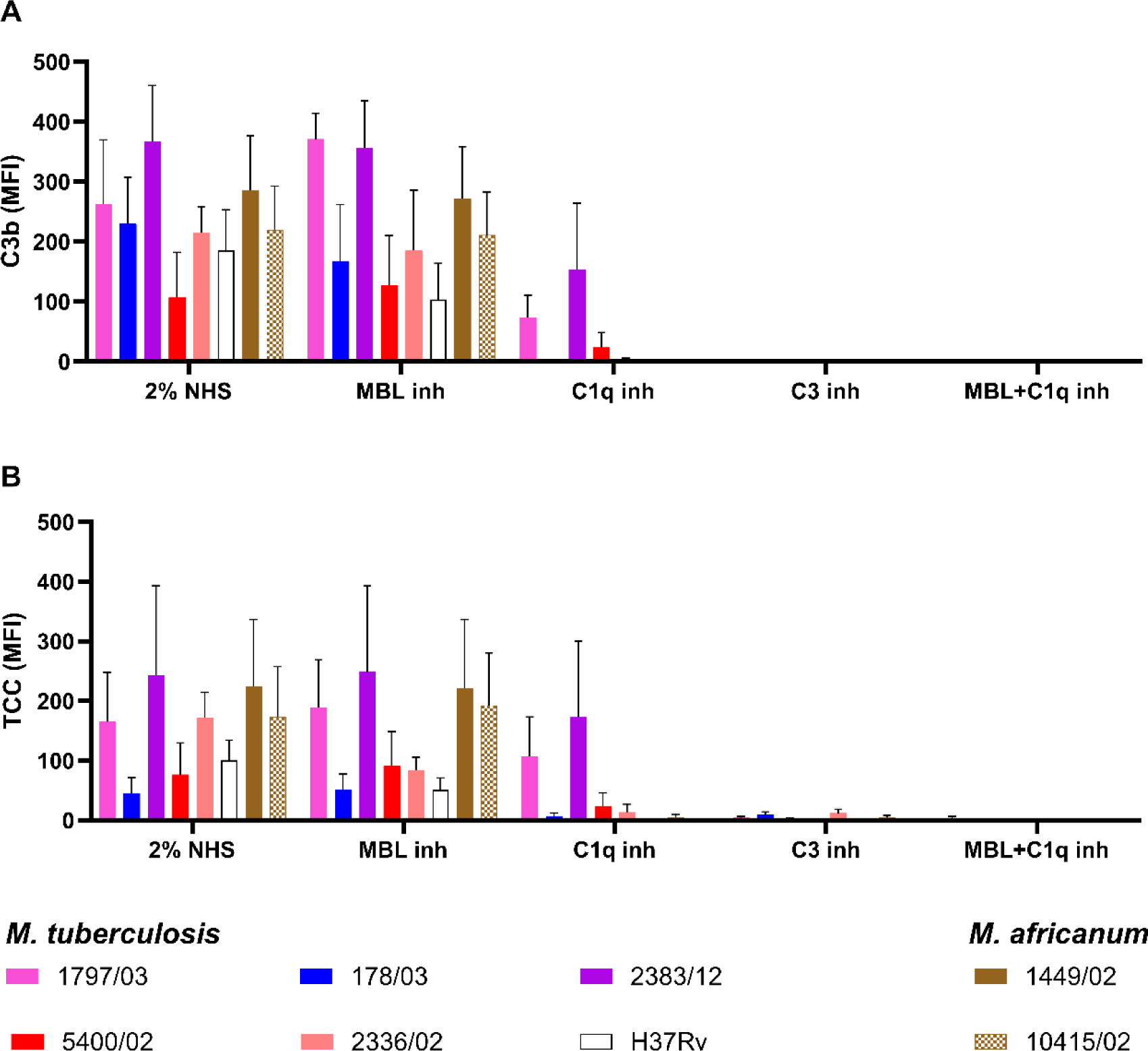
Complement activation on different *Mycobacterium tuberculosis* complex strains in 2% serum. Strains of 6 MTBC lineages were irradiated and incubated with 2% NHS which was pre-incubated with the different inhibitors and mock controls for 10 min at room temperature. Deposition of C3b (**A**) and TCC (**B**) was measured by flow cytometry after incubation with specific mAb followed by the addition of fluorophore-labelled secondary antibodies. The MFI of positive cells was analyzed from at least three independent experiments, negative controls were acquired in the absence of 2% NHS and subsequently subtracted in the analysis. Results represent the means of at least three independent experiments ± SEM.

### Complement activation by main lineages of the MTBC lineages via the AP

To investigate complement activation with the full contribution from the AP, we applied the same strategy using C1q and MBL inhibitors, but in 10% instead of 2% NHS. The serum was added to the bacteria and subsequently C4b (see **Fig S4)** C3b and TCC (**Fig 5**) deposition was examined. In the presence of 10% NHS, C3b and TCC bound to all strains investigated, which confirms our previous results (**Fig 4**) that all different MTBC lineages activate the complement cascade. Also, similar to our previous experiment, the inhibition of MBL did not affect the deposition of C3b and TCC (**Fig 5**). The simultaneous inhibition of C1q and MBL fully reduced C3b and TCC on 2 of the tested strains (2883/12 and 5400/02). This contrasted with the co-inhibition in 2% NHS where deposition was hampered on all the strains. In 10% NHS, the strains most affected at the C3b level were those displaying the lowest baseline C3b levels (10% NHS, no inhibition) and these were also the strains where co-inhibition reduced C4b the most (**Fig S4**). On strains 1797/03, 2336/02 and 1449/02, the C1q/MBL co-inhibition resulted in an approximate 50% reduction of C4b, yet without reducing C3b levels. This suggests that the remaining activity of the LP and CP after inhibition is enough to stimulate the AP amplification of C3b. The incubation of 10% NHS with the respective mock inhibitors did not affect complement activation (**Fig S4**) and similar to our previous experiment, the C3 inhibitor strongly reduced C3b and TCC deposition (**Fig 5**).

**Figure 5.**
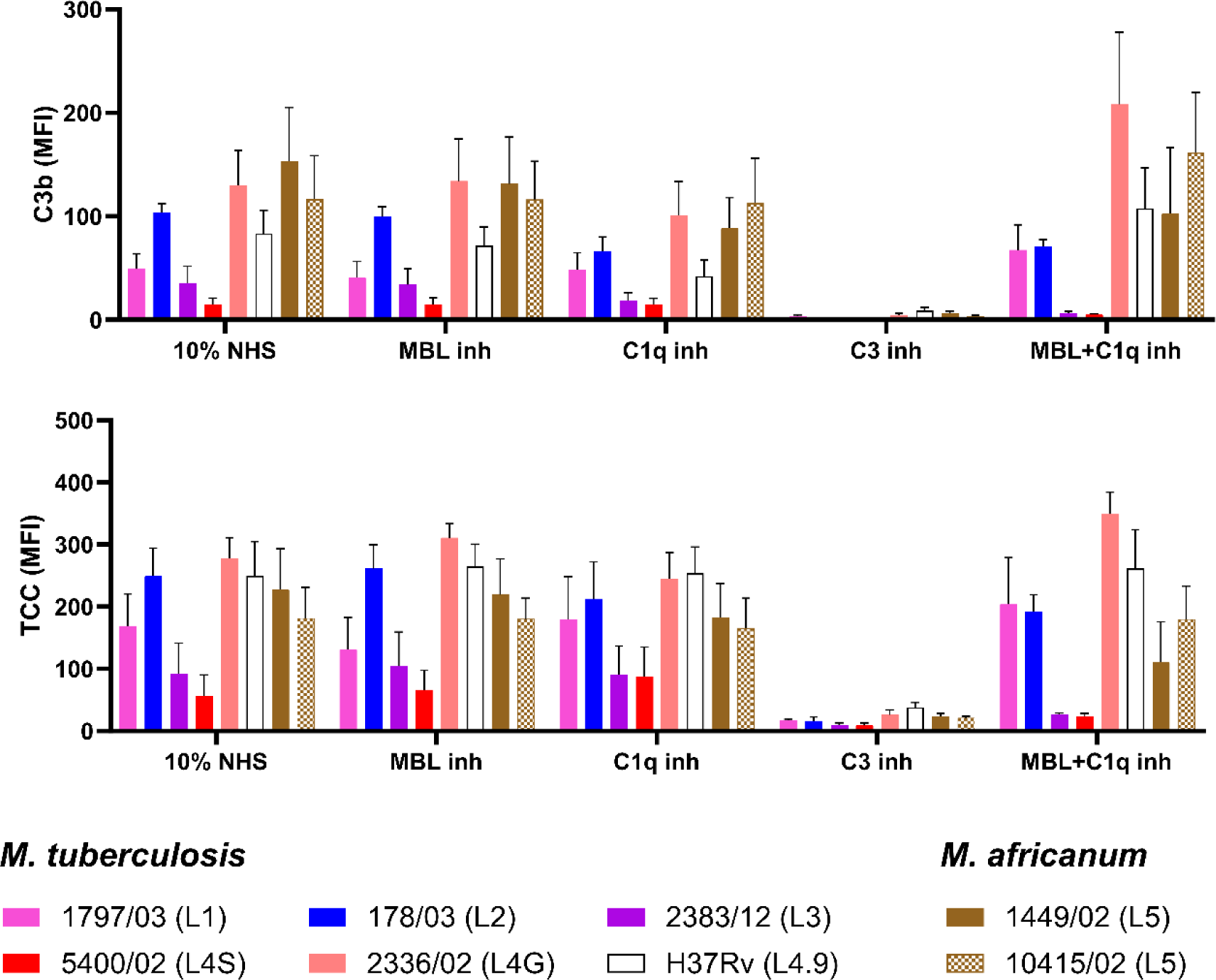
Complement activation on different clinical *Mycobacterium tuberculosis* complex strains in 10% serum. Clinical strains of 6 MTBC lineages were irradiated and incubated with 10% NHS which was pre-incubated with the different inhibitors and mock controls for 10 min at room temperature. Deposition of C3b (**A**) and TCC (**B**) was measured by flow cytometry after incubation with specific mAb followed by the addition of fluorophore-labelled secondary antibodies. The MFI of positive cells was analyzed in triplicates, negative controls were acquired in the absence of 10% NHS and subsequently subtracted in the analysis. Results represent the means of at least three independent experiments ± SEM.

### Impact of serum on survival of mycobacteria

Since we observed deposition of C5b-9 on the surface of MTBC strains, we next investigated whether the presence of NHS affects the viability of the clinical strains. For a comprehensive analysis, viable mycobacteria were incubated with increasing concentrations of NHS generated from cohorts with known BCG vaccination status and, the effect on viability was determined based on subsequent plating and determination of colony-forming units (CFU). The broad range of investigated NHS concentrations tested neither altered the CFU counts of the *M. tuberculosis* strains 1797/03 (L1), 178/03 (L2), 2383/12 (L3), 2336/02 (L4G), 5400/02 (L4S), and H37Rv nor of the *M. africanum* strains 1449/02, 10415/02 (L5) (**Fig 6**), indicating that serum does not affect the viability of MTBC strains.

**Figure 6.**
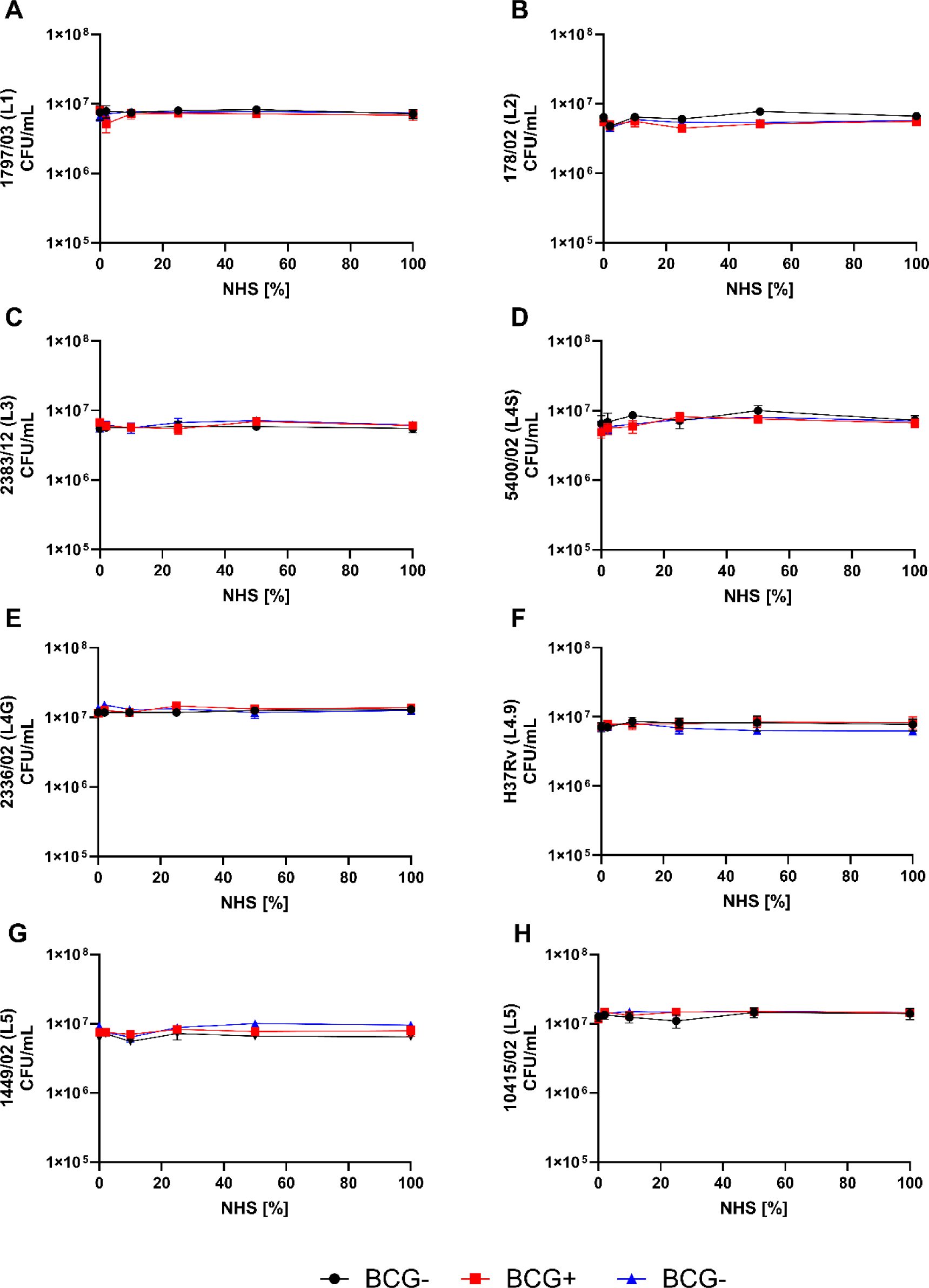
Effect of serum in the viability of *Mycobacterium tuberculosis* complex strains. Clinical strains of 6 MTBC lineages were incubated with 0%, 2%, 10%, 25%, 50% and 100% of NHS from cohorts with different BCG vaccination statuses (BCG-: BCG negative non-vaccinated “serum pool from University Hospital of Copenhagen” (black circle), and BCG+: BCG positive vaccinated (red square) and BCG-: BCG negative non-vaccinated (blue triangle) “serum pool from the Research Center Borstel”), for 30 minutes before plating for subsequent CFU detection. Three independent experiments with technical duplicates were performed and plotted as representing ± SEM.

## Discussion

Although TB is the most prevalent bacterial infectious disease in humans [30] and its pathogenesis is driven by the interplay of pathogen and host factors, the relevance of the complement system, activated by the CP, the LP, or the AP, has not been adequately investigated [31]. In the present study, we provide the first data to understand the interaction between the complement system and a diverse set of clinical MTBC strains representing the global pathogen diversity with a particular focus on strains of globally spreading (generalist), and geographically restricted (specialist) lineages.

Most studies on the interaction of the complement system and mycobacterial strains have been performed using lab-adapted reference strains like H37Rv or *M. bovis* BCG [23, 24]. However, we and others have recently shown that strains of the MTBC have a high pathobiological diversity also impacting mechanisms of the innate immune response that obviously depend on the properties of the bacteria and thus on their genetic makeup [10, 26, 32–34]. To reasonably investigate the sensing of mycobacterial infection by the complement system, we examined the interaction of all complement pathways with representative MTBC strains in this study. To this end, we included 6 lineages represented by 21 strains to provide a comprehensive overview of the relevance of complement in TB. We were able to show the direct binding of MBL and C1q and the importance of C1q for the complement activation on a broad range of clinical MTBC strains.

By investigating the PRMs of the LP using different clinical strains from L1, L2, L3 L4, L5, L6, and the lab-adapted strain H37Rv (which belongs to L4), we found that mainly MBL and to a lesser extent, ficolin-2 can recognize strains of all lineages investigated. Almost all tested strains also bound C1q, yet to a lower extent than MBL. Although not statistically proven, our study additionally suggests, that binding of these molecules depends on the strains investigated.

However, this binding was not lineage specific. Some reports using non-pathogenic mycobacteria and the H37Rv reference strain have indicated that MBL might be considered the key factor for activating the lectin pathway upon recognition of mycobacteria [18, 22]. Additionally, ficolin-3 and to a lesser extent ficolin-1 was suggested to recognize *M. tuberculosis* H37Rv, *M. bovis* BCG, and *M. kansasii* [18].

Various techniques demonstrated that MBL is involved in the recognition of mannosylated lipoarabinomannan (Man-LAM), an important component of the mycobacterial cell wall [35]. MBL not only recognizes *M. tuberculosis* but also other slow-growing mycobacteria such as *M. bovis, M. kansasii, M. gordonae,* and to a lesser extent *M. smegmatis* [18, 35]. Importantly, structural polymorphisms in MBL have been associated with a protective effect against TB caused by *M. africanum/M. bovis* but not against the disease caused by *M. tuberculosis* [26]. As expected, all 21 strains of *M. tuberculosis* and *M. africanum* tested in the present study bound to rMBL. Interestingly, our results differ from the literature [26] which showed a higher MBL binding to strains of L6 compared to strains of L4S. The differences are probably due to the quality of the recombinant proteins used.

Our results corroborate a previous study of the interaction of the complement system with planktonic and biofilm of *M. tuberculosis* [29], showing that C1q can bind mycobacteria also in the absence of immunoglobulins. Moreover, our results reveal for the first time that C1q directly recognizes various MTBC clinical strains. The relevance of C1q in TB research became apparent recently when it was described that the expression of C1q is increased in *M. tuberculosis*-infected *Rhesus macaques* [34] and TB patients suffering from active disease compared to latently infected individuals [35–37]. Therefore, C1q was suggested as a biomarker for active TB. Functionally, in this context, it could be possible that the interaction of C1q with MTBC bacteria might have an impact on the immune response to TB.

Based on our findings that C1q and MBL recognize various strains of all lineages, we have chosen 8 strains from different lineages to further evaluate complement activation. Overall and interestingly, the degree of complement activation (C3b and TCC deposition) appeared to differ between strains. Since our studies primarily identified MBL and C1q to recognize mycobacteria, we further determined the relevance of both molecules for the activation of the complement cascade. Because complement activation in 2% NHS was remarkably reduced by C1q and not by MBL inhibition, and because co-inhibition of C1q and MBL inhibitors completely abrogated complement activation, C1q seems to be the main driver of complement activation with minor contribution of MBL.

We observed a striking variation in the level of complement activation between strains and to our surprise the C3b values were overall lower in 10% than in 2% NHS. This could imply that complement regulators might be operating on the mycobacterial surface in high serum concentration, while the effect might be diminished at low serum concentrations. This observation will be subject to further studies.

Co-inhibition of C1q and MBL in 10% NHS only abolished C3b and TCC on a few strains, which were those mostly affected upstream at the C4b level, likely because of low C4b baseline levels. Hence, strain differences also exist in terms of the degree of LP and CP activation. In the presence of C1q and MBL inhibitors, most of the tested strains still generated sufficient C4b to generate downstream C3b deposition, meaning that the inhibitory effect on C4b did not transcend into reduced C3b and TCC levels. This suggests that the AP amplification loop contributes to the total complement activation on MTBC.

In the present study, C3 inhibition by compstatin revealed a clear reduction of C3b and TCC binding on all MTBC strains investigated. Curiously, we observed an increase in C4b deposition when inhibiting C3b which indicates that decreased C3b on mycobacteria, leaves more space for C4b deposition, a phenomenon recently reported by us on the opportunistic fungus *Aspergillus fumigatus* [36]. We here provide evidence for TCC deposition on a broad spectrum of MTBC strains. The level of TCC binding shows some variation between the strains, again demonstrating that the degree of complement activation varies and differences in the mycobacterial cell wall may affect complement activation and formation of TCC [29]. Although we detected the completely formed C5b-9 complex on the surface of all strains analyzed, the complex formation did not result in lytic activity since different NHS concentrations did not affect the viability of MTBC strains. Interestingly, TCC deposition demonstrated in the present study is supported by the recently observed C5b-9 deposition on the surface of planktonic and biofilm of *M. tuberculosis* [29]. Furthermore, *in vivo* studies using C7-deficient mice point to an only transient role of TCC complement components in the control of Mtb infection [37].

In conclusion, this study sheds light on the importance of evaluating MTBC clinical strains in general and specifically in complement research. By using a diverse group of MTBC strains with a broad range of virulence [38] to analyze complement activation in the context of TB, we can conclude that bacteria from the MTBC complex are in general able to activate the complement system but with high variation between strains. Notably, C1q functions as an innate pattern recognition molecule on the surface of MTBC, indicating a yet unknown role of this complement component in TB. Whether recognition by C1q is of advantage for the bacterium or the host remains to be explored.

## Materials and methods

### Phylogenetic tree for MTBC strains

The aligned sequences of 7476 concatenated SNP positions were produced using MTBseq set up on default parameters and used to calculate a maximum likelihood tree with IQ-TREE v1.6.12 [39], with the best fit-substitution model K3Pu+F tested with ModelFinder ModelFinder [40] and, from 1000 bootstrap trees using ultrafast bootstrap option [41]. The consensus tree was then midpoint-rooted and annotated using FigTree v1.4.4 (http://tree.bio.ed.ac.uk/software/figtree/) and iTOL v6.8.1 [42].

### Cultivation of MTBC strains

Clinical strains of the MTBC were provided by the National Reference Center for Mycobacteria (Borstel, Germany) on Löwenstein/Jensen (L/J) medium. Colonies from L/J media were inoculated in 10 ml 7H9 medium supplemented with 10% oleic acid albumin-dextrose-catalase (OADC), 0.05% Tween 80, and 0.2% glycerol and incubated in 30 ml square medium bottles (Nalgene) at 37°C with shaking. Growth to mid-log phase (OD 0.4-0.6) was monitored by measuring the optical density at 600 nm (OD_600_) every second day (Bio-Tek Synergy). Precultures were transferred into a roller bottle system (Corning) and incubated at 37°C/5 rpm. Twenty ml of fresh medium was gradually added to a final volume of 100 ml (main culture). Log-phase bacterial suspensions (OD_600_, 0.8 to 1) were stored in 1 mL aliquots at −80°C for further experiments. Sterility controls (Ziehl-Neelsen staining, blood agar, Brain Heart Infusion Broth, and LB medium) were performed for all precultures and primary cultures.

### Inactivation of mycobacteria

Mycobacteria were cultured under biosafety level 3 conditions whereas complement assays were conducted in laboratory with lower biosafety level. Therefore, the clinical strains had to be inactivated which was achieved by gamma irradiation. Mycobacteria were irradiated with 2.0 kGy or 2.5 kGy in a BioBeam8000 (Gamma-Service Medical GmbH, Leipzig, Germany) equipped with a ^137^Cs gamma-ray source. Killing of MTBC strains was confirmed by incubating the irradiated samples in the MGIT system (Becton, Dickinson GmbH, Heidelberg, Germany) and plating on solid agar and incubation for at least 3 months at 37°C. Aliquots of the inactivated clinical strains were stored at 4°C until further analyses.

### Proteins of the complement system

We used the following recombinant proteins that were in-house expressed and purified as previously described [43]: rficolin-1, rficolin-2, rficolin-3, rMBL, rMASP-1, rMASP-3 and rCL-11. Purified C1q was purchased from Complement Technology, Inc (A099, CompTech, TX, USA).

### Primary antibodies

For the protein binding assay, we used the following in-house produced monoclonal mouse antibodies (Abs): anti-ficolin-1 mAb FCN106, anti-ficolin-2 mAb FCN219, anti-ficolin-3 mAb FCN334, anti-MASP-1/-3/MAP-1 mAb 8B3 and anti-CL-11 mAb Hyb-15. We applied the following commercial Abs: mouse anti-MBL mAb (HYB 131-1, Bioporto Diagnostics, Gentofte, Denmark), and rabbit anti-C1q pAb (A0136, Dako, Glostrup, Denmark). For complement activation experiments, we used rabbit anti-C4c and -C3c pAbs (0369 and F0201, Dako, Glostrup, Denmark), and mouse anti-TCC mAb clone aE11 (011-01, AntibodyChain, Utrecht, Netherlands).

### Binding of complement proteins to MTBC

Seven different MTBC clinical strains belonging to different lineages and the H37Rv reference strains were screened for binding to various complement system proteins. The inactivated mycobacteria were first washed two times in Barbital-BSA (5 mM barbital sodium, 145 mM NaCl, 2 mM CaCl_2_, 1 mM MgCl_2_, 0.5% BSA) with a centrifugation step in between at 2,500×g for 15 min. Bacteria were then incubated with the different recombinant or purified proteins (5 μg/mL) mentioned above, for 30 min at 37 °C. The cells were washed twice in Barbital-BSA and centrifuged at 2,500×g for 10 min. Protein detection was performed using FCN106 (ficolin-1), FCN219 (ficolin-2), FCN334 (ficolin-3), HYB 131-1 (MBL), Hyb-15 (CL-11), 8B3 (MASP-1/-3/MAP-1) and A0136 (C1q). Finally, each sample was stained with the corresponding secondary Abs goat anti-mouse PE (SouthernBiotech), or sheep anti-rabbit FITC. The median fluorescence intensity (MFI) was measured on a BD FACSCelesta™ and analyzed using the software FCS Express 7 with a defined gating process (**Fig S1**).

### Complement activation on MTBC strains measured by flow cytometry

The activation of the complement system on MTBC strains was examined under various conditions. 1×10^6^ bacteria/mL were incubated in 2% or 10% normal human serum (NHS) (BCG-) diluted in barbital/BSA for 30 min at 37°C, then washed and stained with primary or isotype control antibodies (Abs), followed by FITC-conjugated secondary Abs in these combinations: anti-C4c biotinylated/BV421 Streptavidin; anti-C3c pAb/goat anti-rabbit-FITC pAb; anti-TCC aE11 mAb/goat anti-mouse-PE pAb; rabbit IgG isotype/goat anti-rabbit; and mouse IgG1 isotype/goat anti-mouse-PE. Abs were incubated for 30 min at 4°C. Deposition of C4b, C3b, and TCC on the surface of bacteria was measured on a BD FACSCelesta™ and analyzed using the software FCS Express 7.

### Inhibition of complement activation

Inhibition of MBL and C1q was performed using anti-MBL-inhibitory mAb 3F8 [44], and anti-C1q mAb clone CLB/C1q85 isotype IgG1 (MW1828, Sanquin, Amsterdam, Netherlands). For mock inhibition, anti-MBL mAb 1C10 and irrelevant mouse IgG1κ (557273, BD Biosciences, Albertslund, Denmark) [44]. Before incubation with the bacteria, 2% and 10% of NHS (BCG-strains were incubated in 2% NHS containing the following mAbs and inhibitors: MBL (3F8) (5 μg/ml), C1q85 (5 μg/ml), compstatin (CP40, 6 μM) or mock inhibitor anti-MBL mAb 1C10 (5 μg/ml) and mouse IgG1κ isotype (5 μg/ml). The same procedure was performed using 10% NHS, with the concentration of inhibitors and mock controls doubled.

### Normal human serum killing assay

Frozen aliquots of the different mycobacterial strains were thawed, and approximately 1×10^6^ CFU/mL were washed with Barbital-BSA buffer. After discarding the supernatant, the bacteria were brought into contact with different concentrations of NHS (0%, 2%, 10%, 25%, 50% and 100%). Different types of NHS from different cohorts were used, BCG-not vaccinated (cohort from Copenhagen), and two other groups, BCG+ and BCG-, where the donors were vaccinated and not vaccinated with BCG (cohort from Borstel), respectively. The mixture was then incubated for 2 hours at 37 °C. After this incubation period, serial dilutions were prepared, plated in 7H10 Middlebrook Agar, and incubated at 37 °C until the colonies appeared [45].

## Supporting information

Duque_et_al_2024 Supplement

## Acknowledgements

We thank Alexandra Hölscher, Bettina Eide Holm, Johanna Volz, Doreen Beyer, Silvia Maaß and Mènie Wiemer for the excellent technical assistance. We are grateful to the Core facility of Fluorescence Cytometry at the Research Center Borstel for their support in analyzing flow cytometry data.

## Author contributions

**Conceived and designed the experiments:** PG, SN, AR, SH, CH, KW

**Performed the experiments:** MAD, MPG, ER

**Analysed the data:** MAD, MPG, ER

**Funding acquisition:** CH, SN, PG

**Contributed reagents/materials/analysis tools:** PG, AR, SN, SH, CH

**Supervision:** KW, AR

**Writing – original draft:** MAD, KW

**Writing – review & editing:** MAD, MPG, ER, SH, SN, PG, CH, KW, AR

