## Supplementary material for "C1q in non-immune human serum has a non-redundant complement function towards clinical Mycobacterium tuberculosis complex strains": Duque_et_al_2024 Supplement

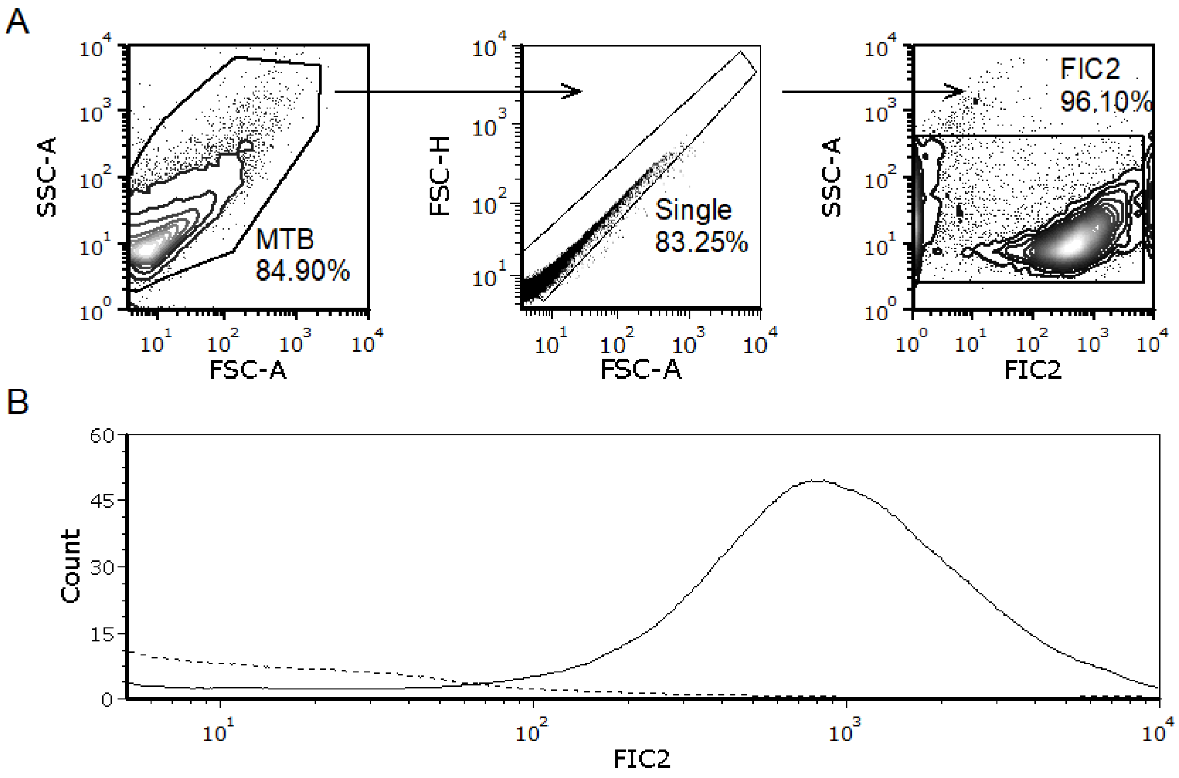


**S1 Fig Gating strategy for detecting the binding of complement PRM to mycobacteria**.

(**A**) The bacterial population was detected by using voltages of 330v for side scattering (SSC) and forward scattering (FSC)**.** Single cells were sub-gated by FCS-H / FSC-A properties before positive cells were detected based on the corresponding fluorophore**.** (**B**) Representative histograms of samples that were incubated with the recombinant or purified PRM (straight line; here Ficollin-2 as an example), and of negative controls where mycobacteria were processed identically but in the absence of PRM (interrupted line). All the samples were plotted, and the corresponding mean fluorescence intensities were extracted.

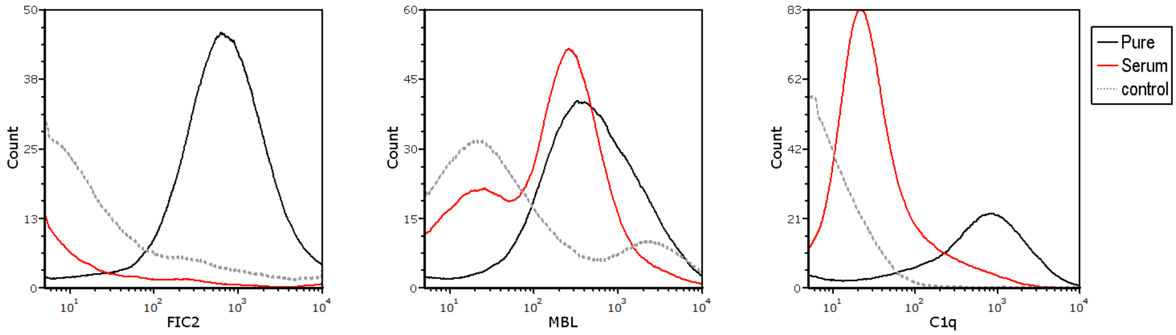


**S2 Fig Comparison of the binding between different PRM using recombinant, pure protein, or NHS.**

The strain 5390/02 was used to compare the binding of, FIC2, MBL and C1q in the presence of pure protein (positive control), NHS (red line) and absence of pure or recombinant protein or serum (grey interrupted line). As the binding in serum for FIC-2 and C1q was low, but we identified C1q binding with pure proteins, we decided to use with all 22 MTBC strains only pure or recombinant proteins.

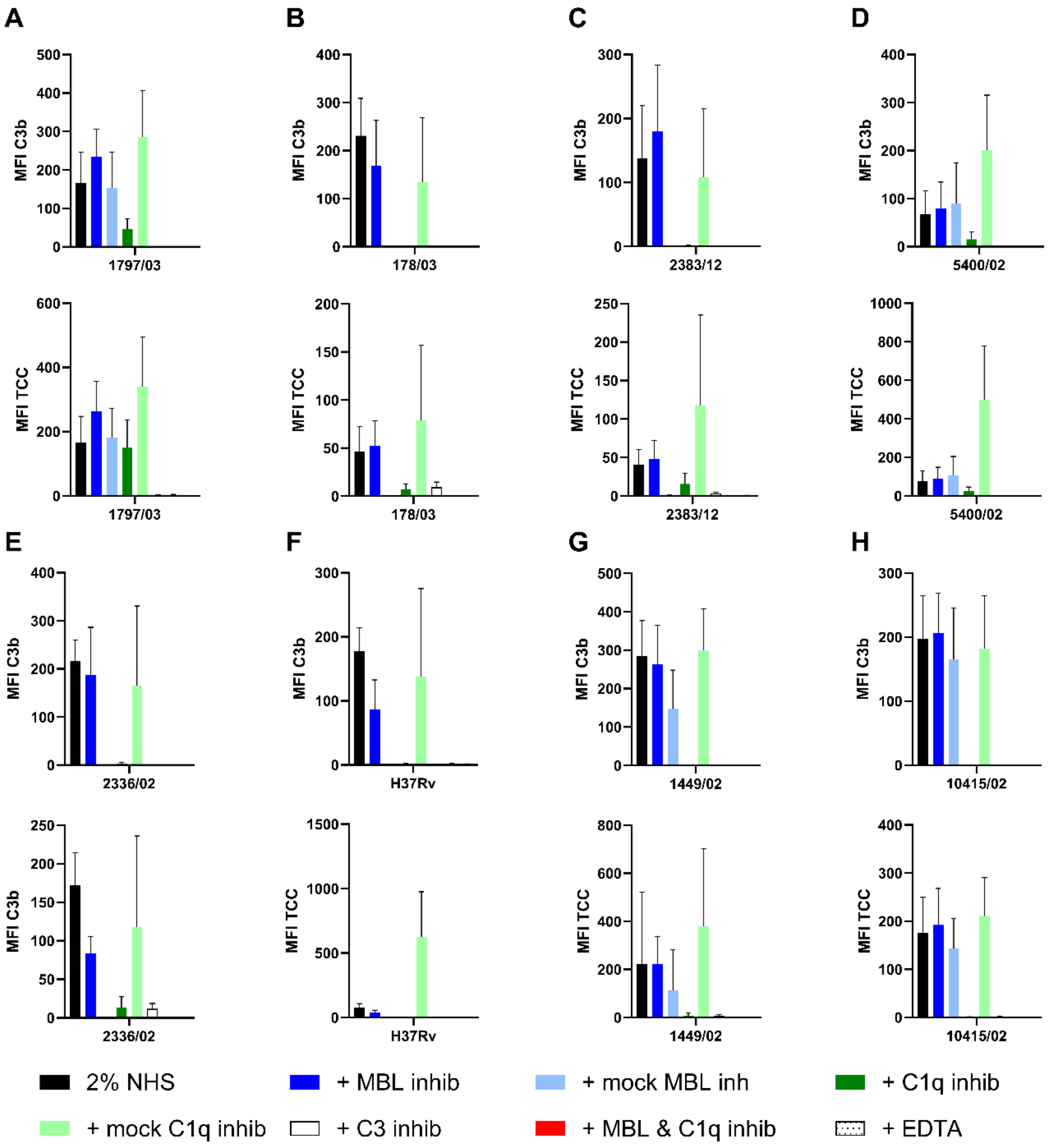


**S3 Fig C3b and TCC deposition on clinical isolates in the presence of 2% NHS.**

1797/03 (L1) (**A**), 178/03 (L2) (**B**), 2383/12 (L3) (**C**), 5400/02 (L4S) (**D**), 2336/02 (L4G) (**E**), H37Rv (L4.9) (**F**) 1449/02 (L5) (**G**), 10415/02 (L5) (**H**). The strains were incubated with 2% NHS with the different inhibitors or mock controls (anti-MBL mAb 1C10 was used as MBL mock-inhibitor and Mouse IgG1 isotype control (BD Biosciences) for C1q mock-inhibitor, for 10 min at room temperature (RT). The MFI of positive cells was analyzed in triplicates, negative controls were acquired in the absence of NHS and subsequently subtracted in the analysis. Results represent the means of at least three independent experiments ± SEM.

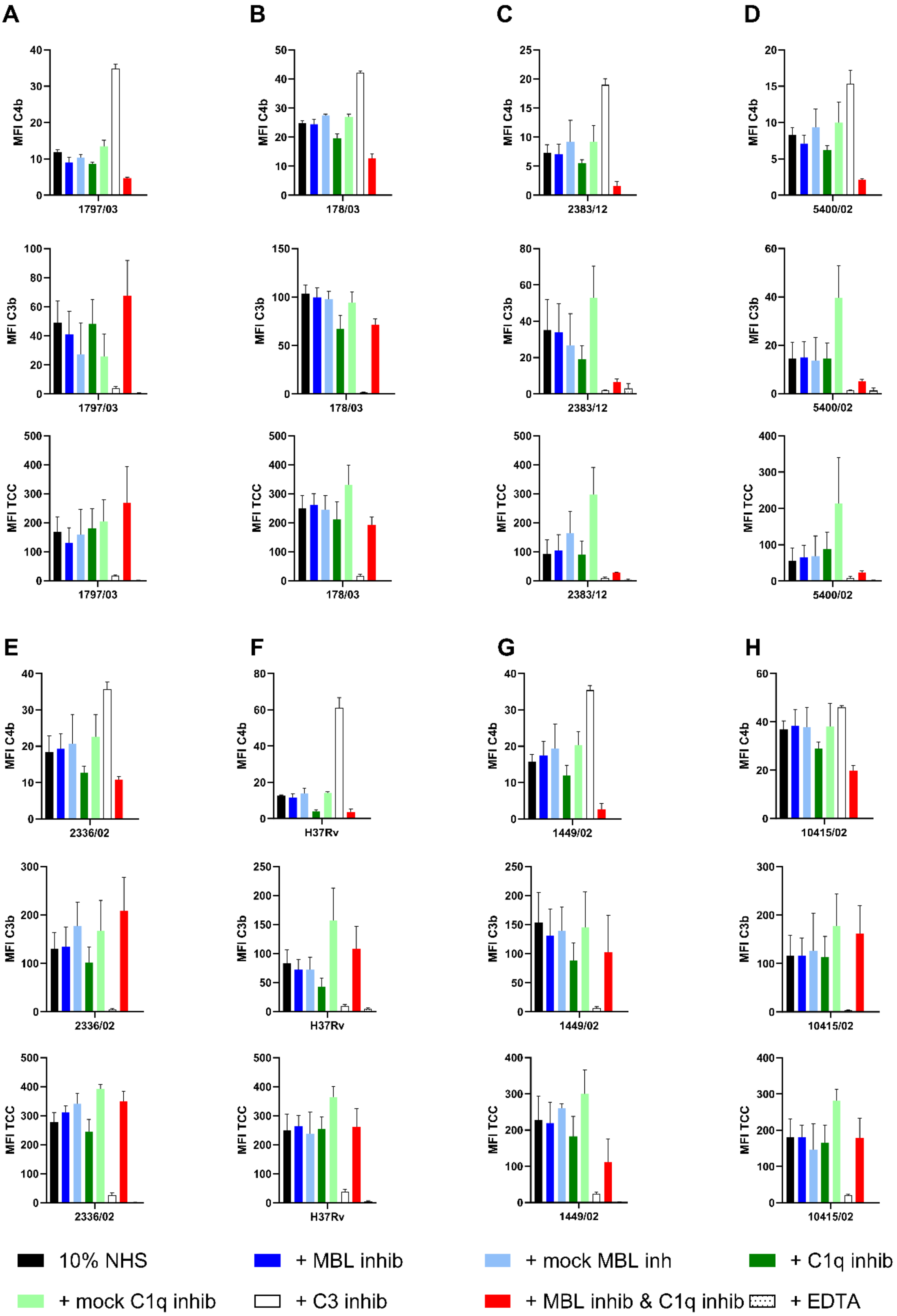


**S4 Fig C4b, C3b and TCC deposition on clinical isolates in the presence of 10% NHS.**

1797/03 (L1) (**A**), 178/03 (L2) (**B**), 2383/12 (L3) (**C**), 5400/02 (L4S) (**D**), 2336/02 (L4G) (**E**), H37Rv (L4.9) (**F**) 1449/02 (L5) (**G**), 10415/02 (L5) (**H**). The strains were incubated with10% NHS with the different inhibitors or mock controls (anti-MBL mAb 1C10 was used as MBL mock-inhibitor and Mouse IgG1 isotype control (BD Biosciences) for C1q mock-inhibitor), for 10 min at room temperature (RT). The MFI of positive cells was analyzed in triplicates, negative controls were acquired in the absence of NHS and subsequently subtracted in the analysis. Results represent the means of at least three independent experiments ± SEM.
